# Awake fMRI Reveals Brain Regions for Novel Word Detection in Dogs

**DOI:** 10.1101/178186

**Authors:** Ashley Prichard, Peter F. Cook, Mark Spivak, Raveena Chhibber, Gregory S. Berns

## Abstract

How do dogs understand human words? At a basic level, understanding would require the discrimination of words from non-words. To determine the mechanisms of such a discrimination, we trained 12 dogs to retrieve two objects based on object names, then probed the neural basis for these auditory discriminations using awake-fMRI. We compared the neural response to these trained words relative to “oddball” pseudowords the dogs had not heard before. Consistent with novelty detection, we found greater activation for pseudowords relative to trained words bilaterally in the parietotemporal cortex. To probe the neural basis for representations of trained words, searchlight multivoxel pattern analysis (MVPA) revealed that a subset of dogs had clusters of informative voxels that discriminated between the two trained words. These clusters included the left temporal cortex and amygdala, left caudate nucleus, and thalamus. These results demonstrate that dogs’ processing of human words utilizes basic processes like novelty detection, and for some dogs, may also include auditory and hedonic representations.

## Introduction

Because dogs can learn basic verbal commands, it is obvious that they have the capacity for discriminative processing of some aspects of human language. For humans, words represent symbolic placeholders for a multitude of people, objects, actions, and other attributes. However, just because a dog can match a word with an action, like ‘fetch,’ does not mean that the dog understands the word has meaning in the same way humans do. For example, dogs may rely on other cues to follow verbal commands such as gaze, gestures and emotional expressions, as well as intonation (Fukuzawa et al., 2005; Mills, 2015; Müller et al., 2015; Persson et al., 2015; D’Aniello et al., 2016). This raises the question of what cognitive mechanisms dogs use to differentiate between words, or even what constitutes a word to a dog.

Part of the problem in studying word comprehension in dogs is the necessity of a behavioral response to demonstrate understanding. Some dogs can retrieve a named object based on a command combined with the name of the object, but this often requires months of training. Examples include Chaser, the border collie who learned over one thousand object-word pairings, and the border collie Rico, who demonstrated the ability to select a novel object among familiar objects based on a novel label (Kaminski et al., 2004; Pilley and Reid, 2011; Zaine et al., 2014).

But these dogs may have been exceptional. Few other dogs have been documented to have this level of expertise. It may be that most dogs rely on simple mechanisms of discrimination – like novelty detection – coupled with other cues from the human to figure out an appropriate behavioral response.

The auditory oddball task, where subjects behaviorally discriminate between target and novel acoustic stimuli, is a well-established task used to measure the processing of target detection and decision-making in humans and nonhumans. The neural regions responsible for target detection and novelty processing not only include primary sensory areas associated with the stimulus modality, but also recruit broader areas such as the posterior cingulate, inferior and middle frontal gyri, superior and middle temporal gyri, amygdala, thalamus, and lateral occipital cortex (Linden et al., 1999; Kiehl et al., 2001; Brazdil et al., 2005; Goldman et al., 2009; Cacciaglia et al., 2015). This suggests that differentiating between target versus novel sounds requires primary auditory cortex as well as an additional attentional network to discriminate between competing sensory stimuli. At least one event-related potential study in dogs suggested similar mechanisms might be at work, finding mismatch negativity to deviant tones (Howell et al., 2012).

Recent advances in awake neuroimaging in dogs have provided a means to investigate many aspects of canine cognition using approaches similar to those in humans. Since 2012, pet dogs have been trained using positive reinforcement to lie still during fMRI scans in order to explore a variety of aspects of canine cognition (Berns et al., 2012; Berns et al., 2013). These studies have furthered our understanding of the dog’s neural response to expected reward, identified specialized areas in the dog brain for processing faces, observed olfactory responses to human and dog odors, and linked prefrontal function to inhibitory control (Cook et al., 2014; Berns et al., 2015; Dilks et al., 2015; Cook et al., 2016a; Cuaya et al., 2016). In one fMRI study, dogs listened to human and dog vocalizations through headphones and showed differential activation within regions of the temporal and parietal cortex (Andics et al., 2014). A follow-up study suggested a hemispheric bias for praise words versus neutral words, a finding that was interpreted as proof of semantic processing in dogs. However, a subsequent correction in which left and right were reversed raised questions about the interpretability of this finding (Andics et al., 2016).

To examine auditory processing in dogs, we used fMRI to measure activity in dogs’ brains in response to both trained words and novel pseudowords. Over several months prior to scanning, owners trained their dogs to select two objects based on the objects’ names. During the fMRI session, the owner spoke the names of the trained objects as well as novel pseudowords the dog had never heard before. If dogs discriminate target words from novel words as humans do, they should show differential activity in the parietal and temporal cortex in response to trained words relative to pseudowords (Friederici et al., 2000; Humphries et al., 2006; Raettig and Kotz, 2008). In addition, if dogs use hedonic mechanisms to associate reward value with trained words, then differential activity should also be observed in the caudate.

## Methods

### Ethics Statement

This study was performed in accordance with the recommendations in the Guide for the Care and Use of Laboratory Animals of the National Institutes of Health. The study was approved by the Emory University IACUC (Protocol DAR-2002879-091817BA), and all owners gave written consent for their dog’s participation in the study.

### Participants

Participants were pet dogs from the Atlanta community volunteered by their owners for fMRI training and experiments (Table 1). All dogs had previously completed one or more scans for the project and had demonstrated the ability to remain still during training and scanning (Berns et al., 2012).

**Table 1.** Dogs and their object names.

| Dog | Breed | Age | Sex | Years with fMRI project | Object 1 | Object 2 |
| --- | --- | --- | --- | --- | --- | --- |
| Caylin | Border Collie | 8 | Spayed F | 4 | Monkey | Blue |
| Eddie | Golden Retriever-Lab mix | 6 | Neutered M | 2 | Piggy | Monkey |
| Kady | Golden Retriever-Lab mix | 7 | Spayed F | 4 | Taffy | Yellow |
| Libby | Pit mix | 11 | Spayed F | 4 | Duck | Hedge Hog |
| Ninja | Australian Cattle dog- mix | 2 | Spayed F | 1 | Block | Monkey |
| Ohana | Golden Retriever | 7 | Spayed F | 3 | Blue | Star |
| Pearl | Golden Retriever | 7 | Spayed F | 3 | Duck | Elephant |
| Stella | Bouvier | 6 | Spayed F | 3 | Stick | Tuxy |
| Truffles | Pointer mix | 12 | Spayed F | 2 | Pig | Blue |
| Velcro | Viszla | 8 | Intact M | 3 | Rhino | Beach Ball |
| Zen | Golden Retriever- Lab mix | 8 | Neutered M | 4 | Teddy | Duck |
| Zula | Lab-Mastiff mix | 4 | Spayed F | 1 | Goldie | Bluebell |

Dog's names, breed, age in years when undergoing scanning, sex, years participating in fMRI experiments, and training objects (S+) are listed

### Word-Object Training

In the current experiment, dogs were trained to reliably fetch or select a trained object given the matching verbal name for the object. The dogs were trained by implementing the “Chaser Protocol” in which object names were used as verbal referents to retrieve a specific object (Pilley and Reid, 2011). To keep the task simple, each dog had a set of two objects, selected by the owner from home or from dog toys provided by the experimenters. One object had a soft texture, such as a stuffed animal, whereas the other was of a different texture such as rubber or squeaked, to facilitate discrimination (Fig. 1).

**Fig 1.**
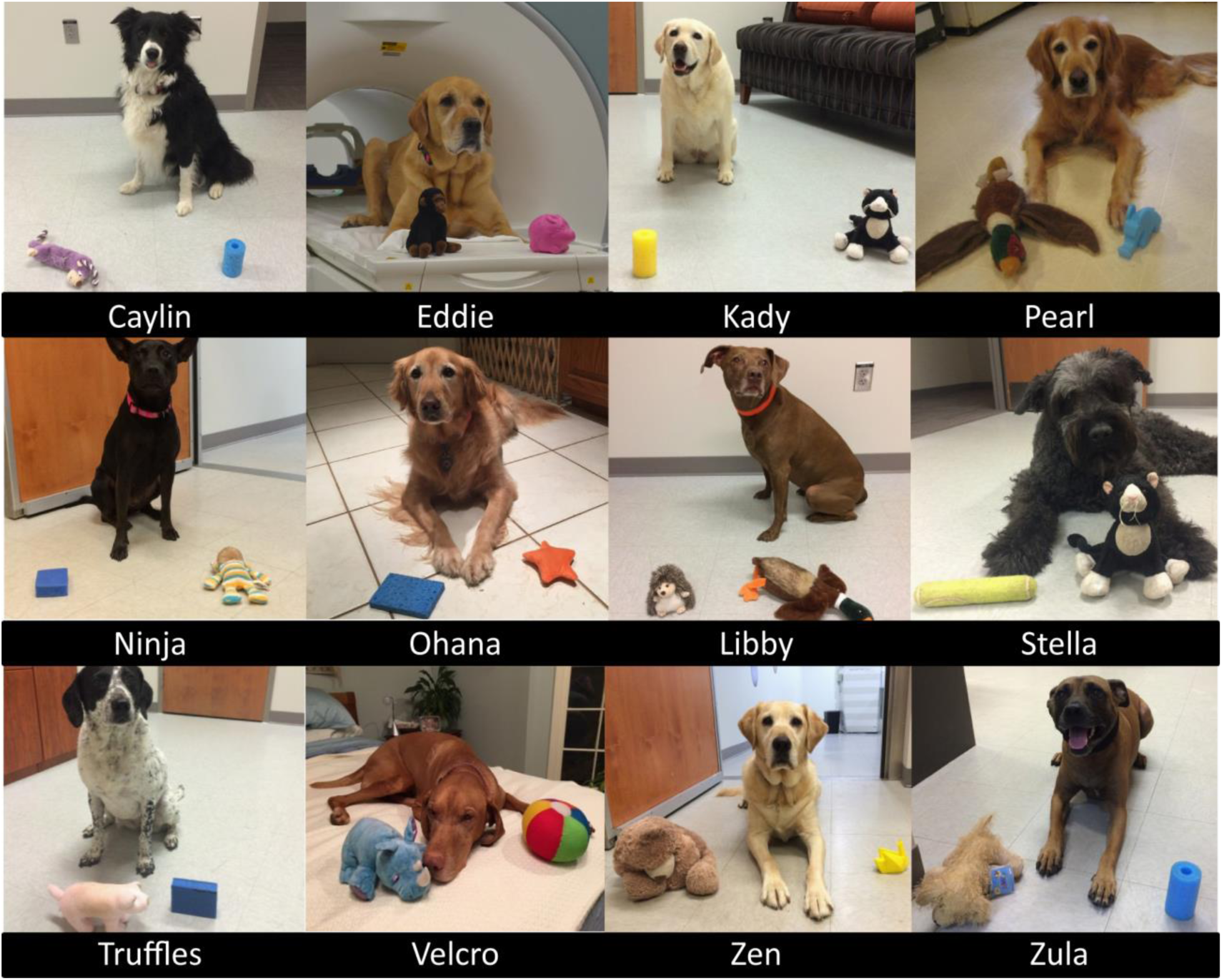
Individual dogs and their trained objects. All 12 dogs successfully trained to retrieve two objects using object names as verbal referents.

Each dog was trained by his or her owner at home, approximately 10 minutes per day, over 2 to 6 months, as well as at biweekly practices located at a dog training facility. Initial shaping involved the owner playing “tug” or “fetch” with her dog and one object while verbally reinforcing the name of the object. Later, the objects were placed at a distance (four feet on average) and the owner instructed the dog to “go get [object]” or “where is [object]?” or “[object]!” The dog was reinforced with food or praise (varied per dog) for retrieving or nosing the object. Next, the object was placed beside a novel object roughly two feet apart, at least 4 feet from the dog, and the command repeated. The dog was reinforced only for correctly selecting the trained object if it was her first selection. Otherwise, if the dog selected the wrong object, the owner made no remark and a new trial began. Regardless of the selection, objects were rearranged before each trial to limit learning by position. If the dog failed to approach an object, the trial was repeated. This training was repeated for each dog’s second object against a different comparison object, to limit the possibility of learning by exclusion. Owners were instructed to train one object per day, alternating between objects every other day until they showed the ability to discriminate between the trained and novel object, at which point they progressed to discrimination training between the 2 trained objects.

All dogs in the current study participated in training for previous fMRI experiments. As described in previous experiments (Berns et al., 2012; Berns et al., 2013; Cook et al., 2014; Cook et al., 2016b), each dog had participated in a training program involving behavior shaping, desensitization, habituation and behavior chaining to prepare for the loud noise and physical confines of the MRI bore inherent in fMRI studies.

### Word-Object Discrimination Tests

Two weeks after progressing to two-object discrimination training, and every two weeks thereafter, each dog was tested on her ability to discriminate between the two trained objects. Discrimination between the two named objects was chosen as the measure of performance, as both objects had a similar history of reinforcement, and this precluded the possibility that performance was based on familiarity. Discrimination testing consisted of the observer placing both trained objects 2-3 feet apart, and at least 4 feet from the dog (Ann Young, 1991), though the number of distractor objects was sometimes increased during training to maximize discriminatory performance. With the dog positioned next to the owner in the heel position, the owner gave the dog the command to “go get [object]” or “[object]!” The dog was reinforced only for correctly selecting the trained object if it was her first selection. If the dog selected the incorrect object, the owner made no remark. After each trial, the objects were rearranged, and the test progressed to the next trial. A performance criterion to move forward to the MRI scan was set at 80% correct for at least one of the objects, with the other object at or above 50%.

During training, owners were asked to report if their dog showed a preference for one object over the other. For the majority of the dogs, the preference was for the softer object of the two, and both the preferred word and the object were consistently labeled as word 1 and object 1. Though Zula passed the discrimination test, she was unable to complete the MRI scan and was excluded from the remainder of the study. Individuals varied on the amount of time needed to train both objects.

### Scan Day Discrimination Test

Scan day tests were conducted in a neighboring room to the MRI room, and were typically conducted prior to the MRI scan. Test procedure was identical to the word-object discrimination test as described above, although the number of trials was increased from 10 to 12 trials if the dog failed to make a response during one or more trials.

### fMRI Stimuli

The stimuli consisted of the two trained words and the corresponding objects. Pseudowords were included as a control condition. Pseudowords were matched to the group of trained words based on the number of syllables and bigram frequency where possible using a pseudoword generator (Keuleers and Brysbaert, 2010) (Table 2). Phoneme substitution was necessary in some cases to ensure that trained words and pseudowords did not overlap at onset or coda. During the scan, pseudowords were followed by the presentation of novel objects with which the dogs had no previous experience. The novel objects included a bubble wand, Barbie doll, stuffed caterpillar, wooden train whistle, plastic gumball dispenser, yellow hat, watermelon seat cushion, Nerf ball launcher, etc.

**Table 2.**
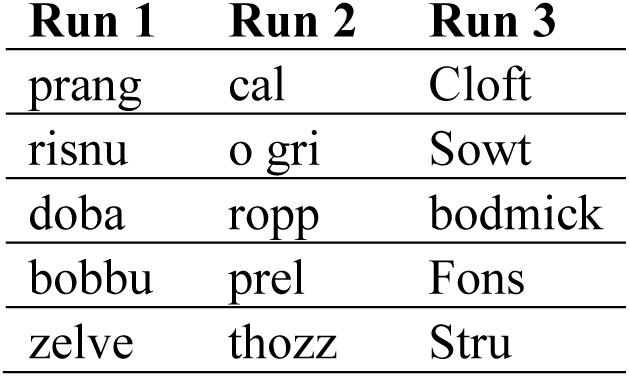
List of pseudowords per run.

| **Run 1** | **Run 2** | **Run 3** |
| --- | --- | --- |
| prang | cal | Cloft |
| risnu | o gri | Sowt |
| doba | ropp | bodmick |
| bobbu | prel | Fons |
| zelve | thozz | Stru |

### fMRI Experimental Design

As in previous studies, dogs were stationed in the magnet bore using custom chin rests. All words were spoken by the dog’s primary owner, who stood directly in front of the dog at the opening of the magnet bore. Both owners and dogs wore ear plugs, which reduced scanner noise by approximately 30 decibels, but allowed for intelligible human speech over the sound of the scanner. The spoken words were intelligible to the experimenters, who also wore ear plugs while next to the MRI during scanning, as well as human operators in the control room via the intercom. At the onset of each trial, a word was projected onto the surface of the scanner, directly above the owner’s head. An experimenter stood next to the owner, out of view of the dog. The experimenter controlled the timing and presentation of the words to the owner via a four-button MRI-compatible button box (Fig. 2A). Onset of words and objects were controlled by the simultaneous presentation and press of the button box by the experimenter marking the onset and duration of presentation. This was controlled manually by the experimenter during each dog’s scan, as opposed to a scripted presentation as in human fMRI studies, because dogs may leave the MRI at any time and data for absentee trials would be lost.

**Fig 2.**
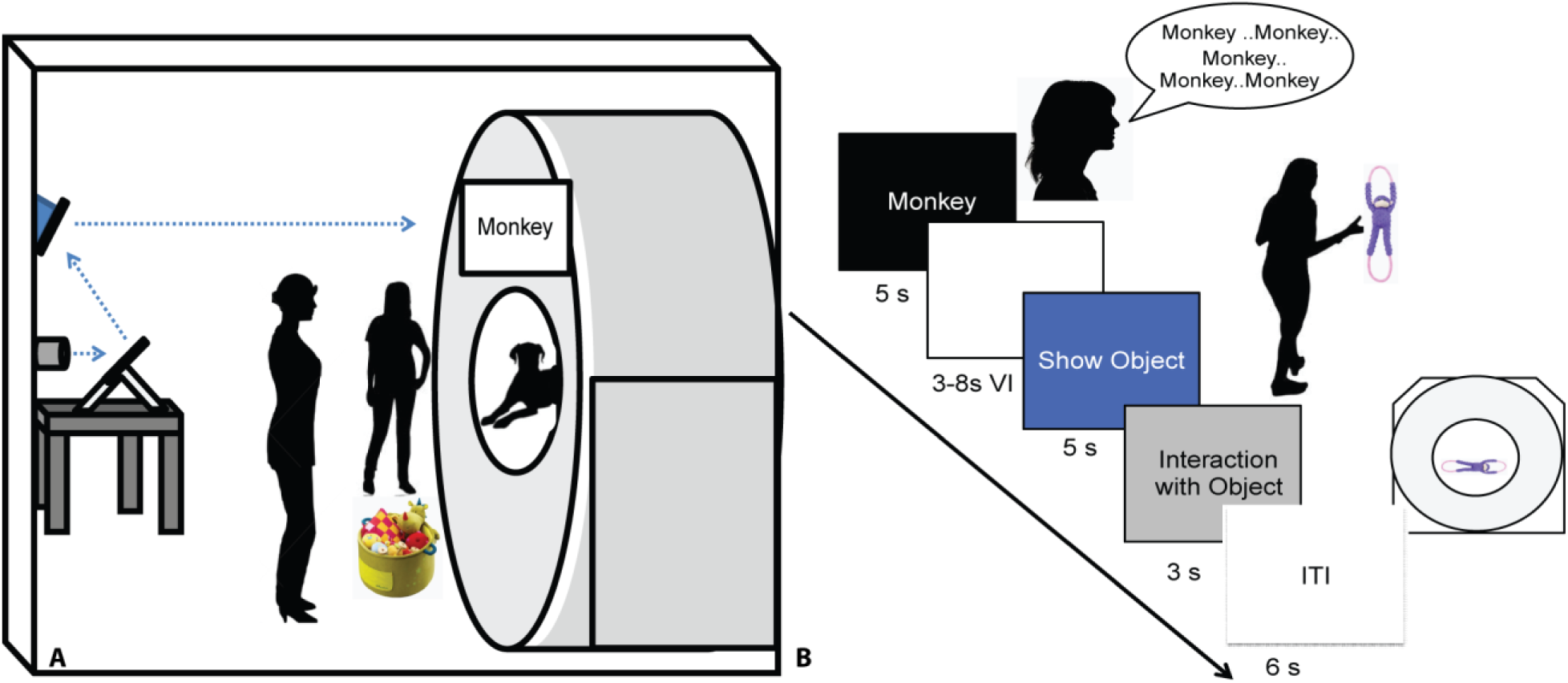
Experimental design. A) Experimental setup with mirror relay projected words onto MRI surface. Owner is facing the projected word and her dog while the experimenter controls the presentation of words and objects to the owner. **B)** Trial timeline indicating spoken word over 5 s, 3-8 s delay, 5 s presentation of object, 3 s for the dog to interact with the object, followed by a 6 s intertrial interval.

An event-based design was used, consisting of four trial types presented semi-randomly: expected, unexpected, pseudoword, and reward. On expected trials, the owner repeated a trained object’s name five times, once per second. Words were repeated to ensure a robust hemodynamic response on each trial and spoken loudly to be heard above the scanner noise. After a variable 3 to 8 s delay, the dog was shown the corresponding object for 5 s and was subsequently allowed to interact with the object. During unexpected trials, the owner repeated the name for a trained object as above, but following the delay period a novel object was presented instead of the corresponding object. In pseudoword trials, the owner repeated a pseudoword, and the delay was followed by a novel object. Reward trials were interspersed throughout each run, during which the owner rewarded the dog’s continued down-stay with food. Trials were separated by a 6 s inter-trial interval, and each dog received the same trial sequence (Fig. 2B). Each of three runs consisted of 26 trials, for a total of 78 trials. The trial types included: 30 expected (15 each of word1 and word2), 15 unexpected (7 or 8 of word1 and word2), 15 pseudowords, and 18 food rewards.

### Imaging

Scanning for the current experiment was conducted with a Siemens 3 T Trio whole-body scanner using procedures described previously (Berns et al., 2012; Berns et al., 2013). During previous experiments, a T2-weighted structural image of the whole brain was acquired using a turbo spin-echo sequence (25-36 2mm slices, TR = 3940 ms, TE = 8.9 ms, flip angle = 131°, 26 echo trains, 128 × 128 matrix, FOV = 192 mm). The functional scans used a single-shot echo-planar imaging (EPI) sequence to acquire volumes of 22 sequential 2.5 mm slices with a 20% gap (TE = 25 ms, TR = 1200 ms, flip angle = 70°, 64 × 64 matrix, 3 mm in-plane voxel size, FOV = 192 mm). Slices were oriented dorsally to the dog’s brain (coronal to the magnet, as in the sphinx position the dogs’ heads were positioned 90 degrees from the prone human orientation) with the phase-encoding direction right-to-left. Sequential slices were used to minimize between-plane offsets from participant movement, while the 20% slice gap minimized the “crosstalk” that can occur with sequential scan sequences. Three runs of up to 700 functional volumes were acquired for each participant, with each run lasting 10 to 14 minutes.

## Analysis

### Preprocessing

Data preprocessing included motion correction, censoring and normalization using AFNI (NIH) and its associated functions. Two-pass, six-parameter affine motion correction was used with a hand-selected reference volume for each dog that best reflected their average position within the scanner. All volumes were aligned to the reference volume. Aggressive censoring (i.e., removing bad volumes from the fMRI time sequence) was used because dogs can move between trials, when interacting with the object, and when consuming rewards. Data were censored when estimated motion was greater than 1 mm displacement scan-to-scan and based on outlier voxel signal intensities. Smoothing, normalization, and motion correction parameters were identical to those described previously (Cook et al., 2016b). A high-resolution canine brain atlas was used as the template space for individual spatial transformations (Datta et al., 2012). The atlas resolution was 1 mm × 1 mm × 1 mm. Thus voxel volumes are in mm^3^.

### General Linear Model

For *a priori* hypotheses, each participant’s motion-corrected, censored, smoothed images were analyzed with a general linear model (GLM) for each voxel in the brain using 3dDeconvolve (part of the AFNI suite). Nuisance regressors included motion time courses generated through motion correction, constant, linear, quadratic, and cubic drift terms. The drift terms were included for each run to account for baseline shifts between runs as well as slow drifts unrelated to the experiment. Task related regressors included: (1) spoken word1; (2) spoken word2; (3) spoken pseudowords; (4) presentation of object1; (5) presentation of object2; (6) presentation of unexpected objects (novel object following either word1 or word2); and (7) presentation of novel objects following a pseudoword. The object on which each dog performed best during the day of the MRI scan as well as the object owners reported as being the preferred of the two was labeled as word1 and object1 when creating the GLM regressors. Stimulus onset and duration were modeled using the dmUBLOCK function, with the 5 utterances treated as a block.

### Whole Brain Analysis

Contrasts focused on the dogs’ response to words and pseudowords. Auditory novelty detection was probed with the contrast: [pseudowords – (word1 + word2)/2]. Low-level aspects of language processing (including acoustic and hedonic representations) were probed with the contrast [word1 – word2] and expectation violation with [novel objects – unexpected objects].

Each participant’s individual-level contrast from the GLM was normalized to template space as described in (Berns et al., 2012; Cook et al., 2014) via the Advanced Normalization Tools (ANTs) software (Avants et al., 2011). Spatial transformations included a rigid-body mean EPI to structural image, affine structural to template, and diffeomorphic structural to template. These spatial transformations were concatenated and applied to individual contrasts from the GLM to compute group level statistics. 3dttest++, part of the AFNI suite, was used to compute a t-test across dogs against the null hypothesis that each voxel had a mean value of zero. All contrasts mentioned above as part of the GLM were included.

As there is spatial heterogeneity within fMRI data, the average smoothness of the residuals from each dog’s time series regression model was calculated using AFNI’s non-Gaussian spatial autocorrelation function 3dFWHMx –acf. The acf option leads to greatly reduced FPRs clustered around 5 percent across all voxelwise thresholds (Cox et al., 2017). AFNI’s 3dClustsim was then used to estimate the significance of cluster sizes across the whole brain after correcting for familywise error (FWE). Similar to human fMRI studies, a voxel threshold of p ≤ 0.005 was used, and a cluster was considered significant if it exceeded the critical size estimated by 3dClustsim for a FWER ≤ 0.01, using two-sided thresholding and a nearest-neighbor of 1.

### Multivoxel Pattern Analysis (MVPA)

In previous fMRI studies of the oddball task, it was noted that attentional differences occurring trial-by-trial may go undetected in the univariate analysis (Goldman et al., 2009). As an exploratory analysis, we used searchlight MVPA to identify regions potentially involved in the representation of words that were not captured in the univariate analysis. We were primarily interested in the representation of word1 vs. word2.

We used a linear support vector machine (SVM) for a classifier because of its previously demonstrated robust performance (Misaki et al., 2010; Mahmoudi et al., 2012). Unsmoothed volumes were censored for motion and outlier count as in the univariate GLM. We then made a model for the unsmoothed data using AFNI’s 3dDeconvolve stim_times_IM function. This model yielded trial-by-trial estimates (betas) for each repetition of word1 and word2, regardless of which object followed. Although it is common in the human literature to use each scan volume as a data point in MVPA (for training and testing), we have found this approach to be problematic with dogs, who move more than humans, resulting in spurious volumes that should be censored. Estimating the beta for each trial affords an additional level of robustness with less sensitivity to potential outlier volumes due to motion. As an additional check for outliers, masks were drawn of the left and right caudate on each dogs’ T2-weighted structural image. Average beta values were extracted from both the left and right caudate for each trial of word1 and word2. Trials with beta values greater than |3%| were assumed to be non-physiological and were removed prior to MVPA. Finally, these trial-dependent estimates were then used as inputs to a whole-brain searchlight MVPA for each individual dog using PyMVPA2 (Hanke et al., 2009). The classifier was trained on the fMRI dataset for each dog by training on 2 runs and testing on the third using the NFoldPartitioner. We used the Balancer function to retain the same number of trials for word1 and word2 across training and testing for 100 repetitions. For the searchlight, we used a 3-voxel radius sphere. This yielded a map of classification accuracies throughout each dog’s brain.

Given the difficulty in finding significant effects in small datasets using cross-validation and parametric methods, we used a permutation approach outlined by Stelzer et al. (2013) to determine the significance of any cluster of common voxels across dogs (Stelzer et al., 2013; Varoquaux, 2017). Briefly, we permuted the order of attributes—but not their corresponding data—and ran the searchlight in individual space for all dogs. This created a null distribution of accuracies. The mean of these distributions was noted to be very close to 0.5, confirming that the classifiers wasn’t biased or skewed. The cumulative distribution of that an accuracy ≥ 0.63 corresponded to the top 5% of voxels, and this was used as a cut-off threshold for the individual maps. These binarized maps were transformed into template space and the average computed across dogs. The resultant group map represented the locations of potentially informative voxels and served as qualitative representation of the relative consistency versus heterogeneity of word-processing in the dogs’ brains. Somewhat arbitrarily, we only considered locations in which at least two dogs had informative voxels.

## Results

### Scan Day Discrimination Tests

Scans were scheduled as close as possible to the day on which object identification criterion was met (*M* = 9.33 days, *SD* =4.92 days) based on owner availability. On the day of the scheduled MRI scan, each dog was tested on her ability to behaviorally differentiate between the two trained objects out of 5 trials each. With the exception of Eddie, each dog correctly selected object 1 on 80 to 100 percent of the trials [*M*=85.73%, *SE* =3.87%], and object 2 on 60 to 100 percent of the trials [*M*=64.27%, *SE*=5.91%] (Fig. 3). The percent correct performance (subtracting 50 percent for chance levels of responding) on scan days for each object was compared in a mixed-effect linear model and showed that performance was significantly greater than chance [*T*(17.1) = 3.00, *P* = 0.008] and that there was a significant difference in performance between word1 and word2 [*T*(11) = 4.67, *P* < 0.001].

**Fig 3.**
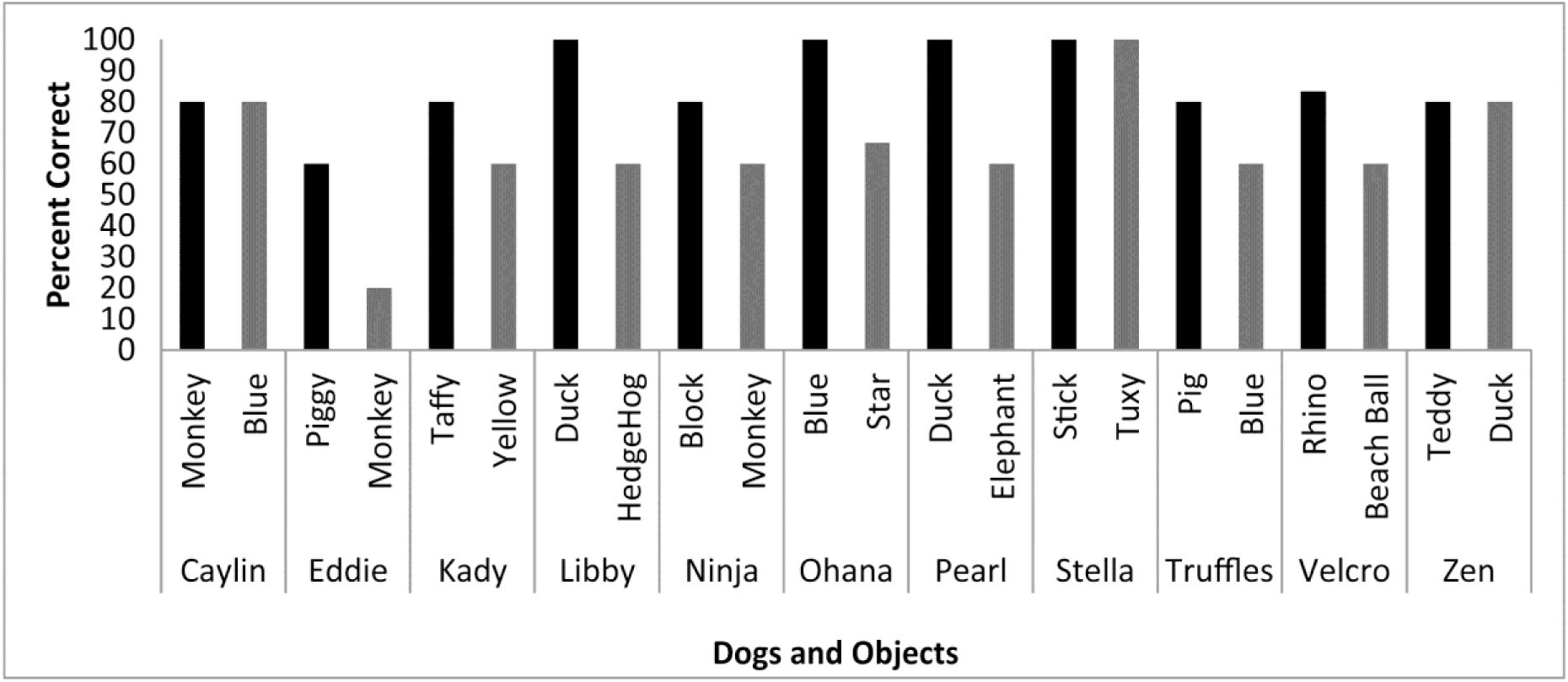
Individual performance on two object discrimination tests. Tests were conducted on the day of the fMRI scan. Each dog’s Object 1 is in black, object 2 is in grey. All dogs performed significantly greater than chance, with the dog’s greater performance or owner’s report of their preference for one object over the other designating object 1.

### Primary Auditory and Visual Activation

To confirm that the dogs clearly heard the words during scanning, a simple contrast subtracting activation to objects (trained and novel) from activation to words (trained and pseudowords) was performed. In human fMRI, the MRI operator may ask the participant whether they can hear auditory stimuli, which is not necessarily possible in dog fMRI, so this was included as a quality check. We opted for an unthresholded image not only to highlight the activation in bilateral auditory cortex but, just as important, to show what was not activated. Notably in the contrast [Words—Objects] positive activation was localized to the auditory cortex for words and negative activation for presentation objects in parietal cortex (Fig. 4), confirming that the dogs heard the words and saw the objects.

**Fig 4.**
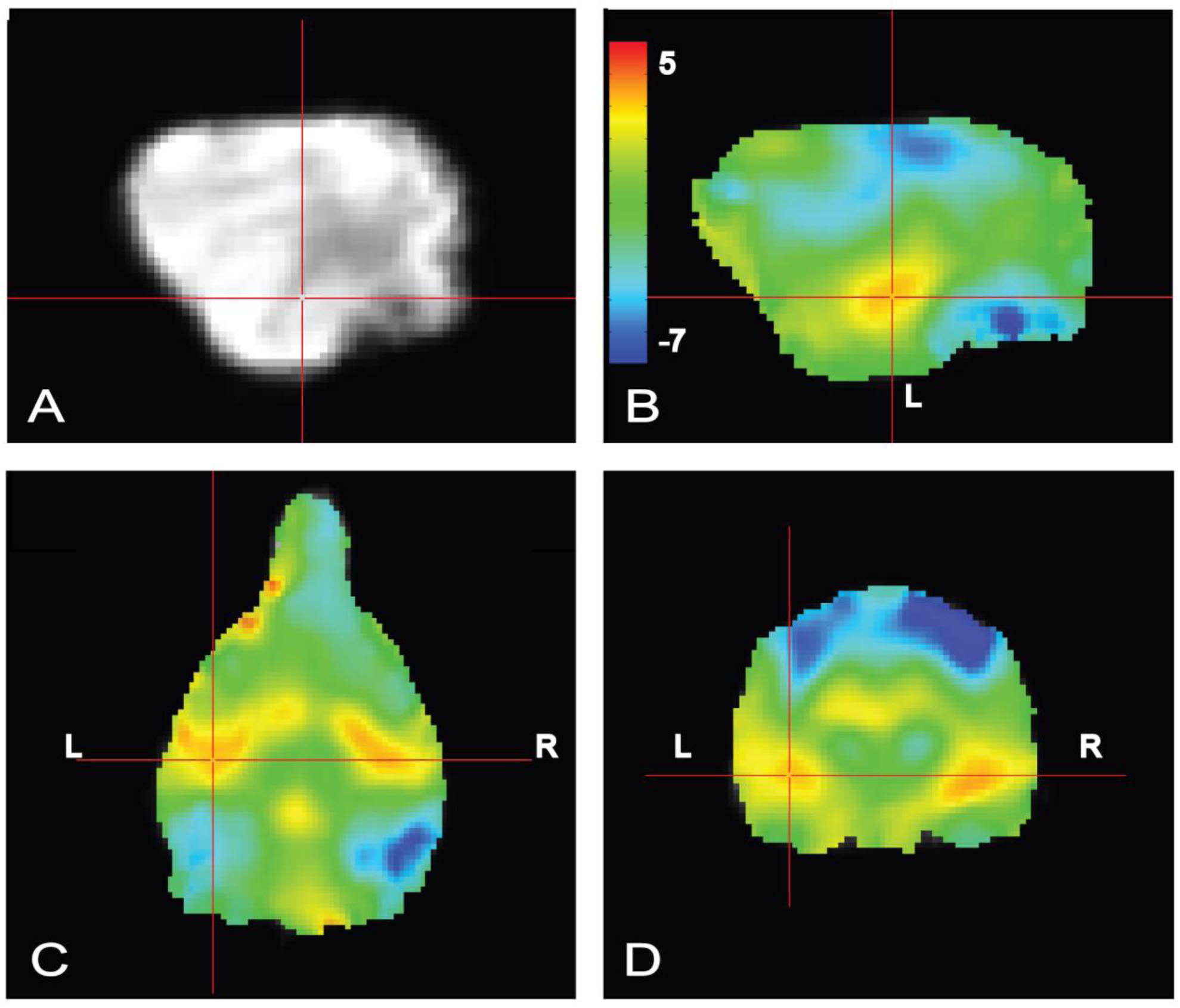
Whole brain group map showing unthresholded activation to all words versus all objects. **A)** Location of crosshairs in superior temporal lobe on average image of all dogs. **B)** Sagittal view of left hemisphere. Colors represent T-statistics. The primary auditory region extending into the parietotemporal area showed greater activation to words (*red*), whereas parietal and occipital areas showed greater activation to objects (*blue*). **C)** Dorsal view. **D)** Transverse view.

### Whole Brain Analyses

Whole brain analysis of the contrasts of interest revealed significant activation only within the right parietotemporal cortex for the contrast [pseudowords – trained words]. With a voxel-level significance threshold of *P* ≤ 0.005, the cluster size in the right hemisphere (839 voxels) was statistically significant at *P* ≤ 0.005 after correction for whole-brain FWE (although activation appeared bilaterally) (Fig. 5). Whole brain analysis of the contrasts of [word1– word2] and [novel – unexpected] were not significant as no cluster survived thresholding at the voxel significance mentioned above.

**Fig 5.**
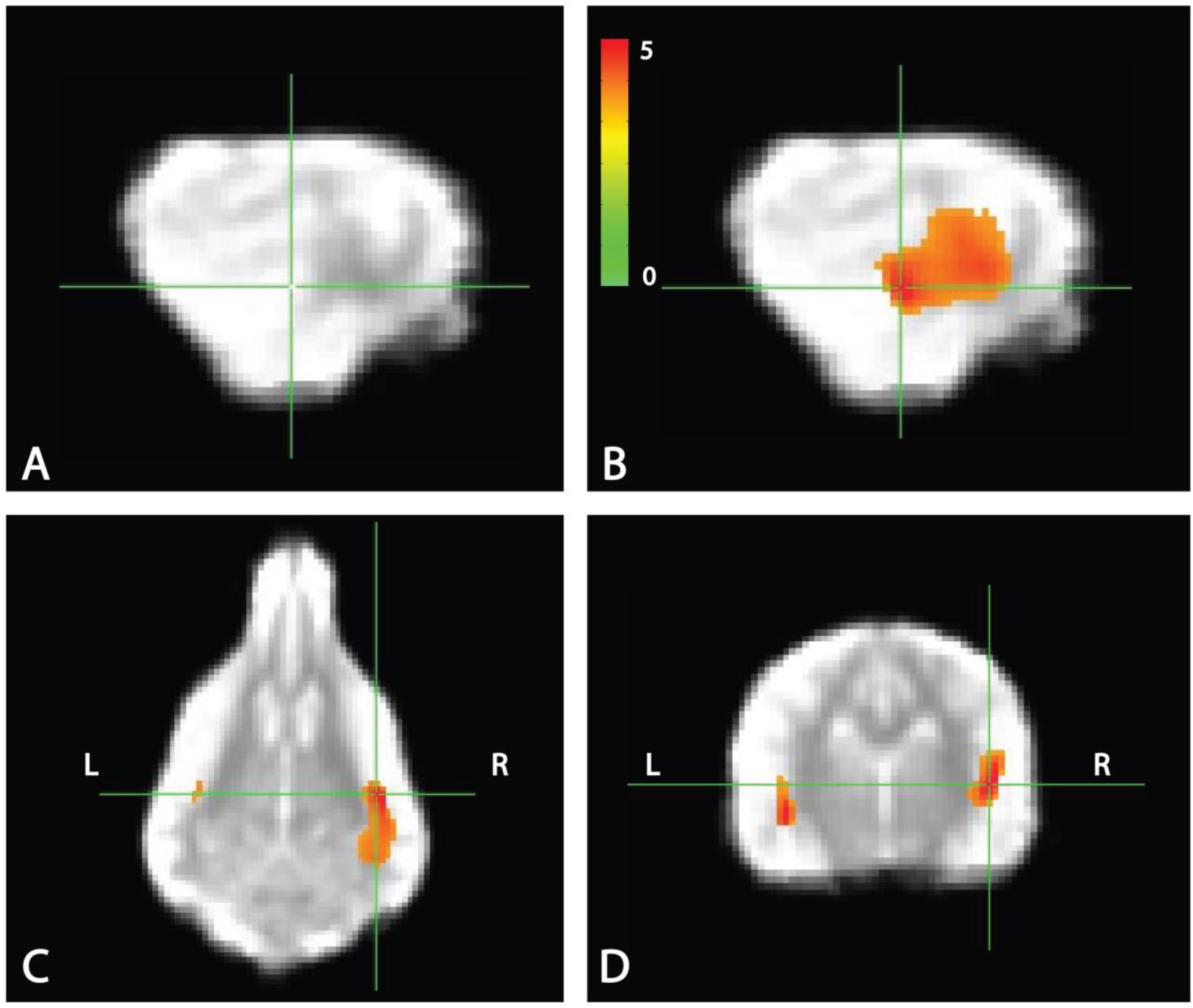
Whole brain response to [pseudowords – words] contrast. Whole brain analysis revealed significant activation within a parietotemporal region including primary auditory cortex and neighboring regions. **A)** Location of crosshairs on average image of all dogs without overlay. **B)** Sagittal view of right hemisphere. Colors represent T-statistics. With a single voxel significance of 0.005, the clusterwise significance (Right: 839 voxels; Left: 43 voxels) corrected across the whole brain was *P* = 0.005 for the right hemisphere, though activation seemed bilateral. **C)** Dorsal view. **D)** Transverse view.

### MVPA

Because the univariate analysis of word1 vs. word2 did not reveal any region with a significant difference, we used MVPA to explore potential regions that may code for different representations of the words. The searchlight map of word1 vs. word2, which identified regions involved in the discrimination of the trained words, showed four clusters of informative voxels (Fig. 6): posterior thalamus/brainstem; amygdala; left temporoparietal junction (TPJ); and left dorsal caudate nucleus. Seven dogs shared informative voxels in or near the left temporal cortex that passed the 0.63 accuracy threshold (Fig. 7).

**Fig 6.**
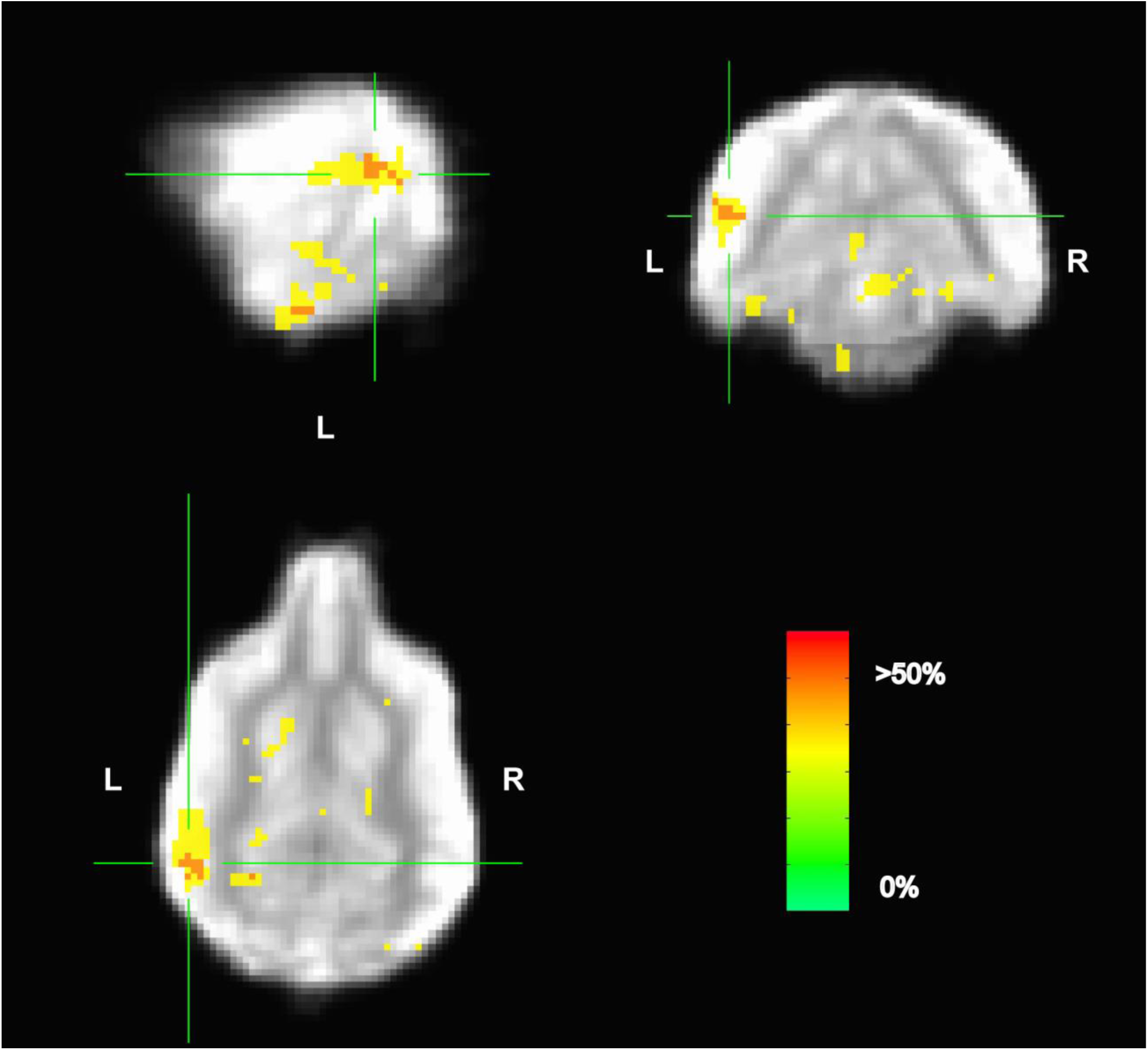
Aggregate performance of searchlight MVPA classifier for word1 and word2 across dogs. Color intensity indicates fraction of dogs with informative voxels at each location. The image is thresholded such that only voxels that were informative for more than one dog are shown. This map showed four clusters: posterior thalamus/brainstem; amygdala; left temporoparietal junction; and left dorsal caudate nucleus. The temporoparietal junction appears similar to human angular gyrus and could be a potential site for receptive language processing in dogs.

**Fig 7.**
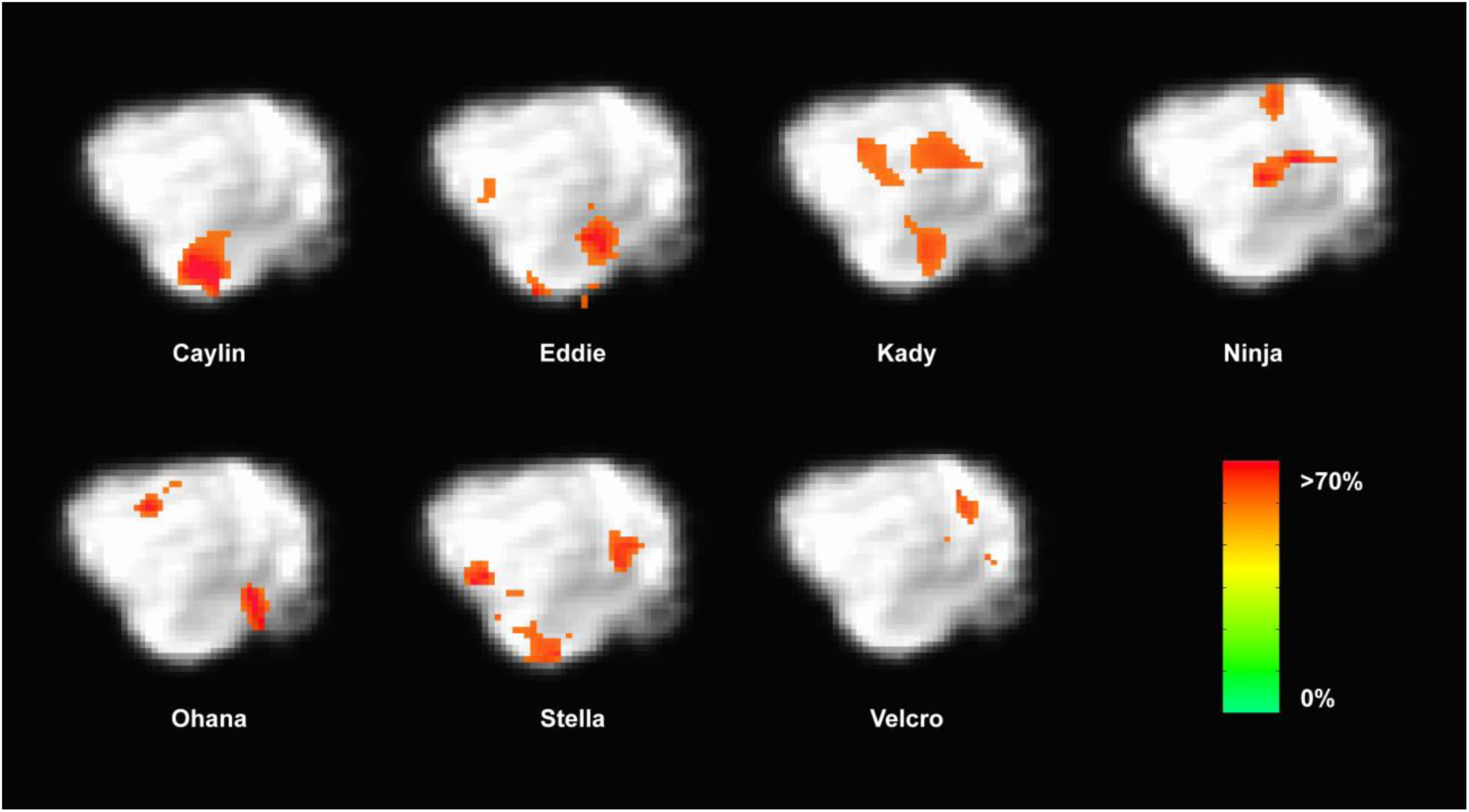
Dogs with informative voxels for word1 and word2 in the left temporal and parietal lobes. Color intensity indicates classification accuracy at each location, thresholded ≥ 0.63. Seven dogs displayed clusters in the left temporal and parietal lobes, suggesting some heterogeneity in the location underlying word discrimination.

## Discussion

Using awake-fMRI in dogs, we found neural evidence for auditory novelty detection in the domain of human speech. The hallmark of this finding was greater activation in parietotemporal cortex to novel pseudowords relative to trained words. Thus, even in the absence of a behavioral response, we demonstrate that dogs process human speech at least to the extent of differentiating words they have heard before from those they have not. The mechanism of such novelty detection may be rooted in either the relatively less frequent presentation of the pseudowords (oddball detection) or the lack of meaning associated with them (lexical processing).

The activation observed in the parietotemporal cortex to pseudowords relative to trained words meets current standards of human fMRI analyses concerning up-to-date methods for cluster thresholds. Specifically, to address concerns raised by Eklund et al. (2016), present analyses for cluster inferences address the former Gaussian-shaped assumption about spatial structure in the residuals of fMRI data and provide more accurate false positive rates compared to previous methods (Eklund et al., 2016; Cox et al., 2017; Slotnick, 2017). As the identified cluster was significant at *P* ≤ 0.005, corrected for whole-brain FWE, the result does not appear to be a false positive. However, as the study was limited to 11 participants, future studies with an increased number of participants could produce a more robust finding.

In humans, greater BOLD response to pseudowords versus real words has been noted in the superior temporal gyrus – an area potentially analogous to the one we identified in dogs (Kotz, 2002; Raettig and Kotz, 2008). In humans, stronger activation to pseudowords depends on whether the pseudoword strongly resembles a known word or is so unlike known words as to prevent any semantic retrieval. When the pseudoword is similar to a known word, more processing has been observed in the superior temporal gyri, presumably to disambiguate it from known words (Raettig and Kotz, 2008). Thus, in dogs, the greater activation to the pseudowords could be due to the acoustic similarity between pseudowords and words that the dogs “knew” and their attempt to resolve the ambiguity. This would be a form of low-level lexical processing. However, previous research has shown that dogs can discriminate between altered phonemes of well-known commands (Fukuzawa et al., 2005), suggesting that it is unlikely that the dogs in our study were confused by acoustic similarity of words and pseudowords.

More likely, a novel word resulted in increased auditory processing to facilitate learning the association with the novel object that followed. A dog’s behavioral bias for novelty is often described as an explanation for performance otherwise labeled as learning by exclusion (Bloom, 2004; Markman and Abelev, 2004; Zaine et al., 2014). As such, a dog may select a novel item because it is novel among other stimuli, but not because she has learned all other stimuli and associated a new word with the novel item. A bias for novelty would therefore be reflected in the dog’s brain as with her behavior.

Auditory stimuli can be difficult to discriminate in the scanner. We used a continuous scanning protocol because that is what the dogs were accustomed to. The simple contrast of all words vs. all objects showed bilateral activation of the superior temporal lobe, indicating that the dogs heard something. However, the main effect of pseudowords vs. trained words showed that the majority of dogs discriminated well enough to tell the difference. The predominant location in the auditory pathway also suggests that the effect wasn’t based on non-verbal cues from the handler (i.e. Clever Hans effect).

The manner in which dogs learn words is different than humans do, and this undoubtedly affects their performance on behavioral tests and the patterns of brain activation we observed. Humans acquire nouns as early as six months of age and differentiate between nouns prior to their ability to use verbs (Bergelson and Swingley, 2012; Waxman et al., 2013). In contrast, dogs do not typically have much experience with nouns because humans tend to train them on actions/verbs (e.g. sit and fetch). Consequently, even the trained words in our study were novel for the dogs in comparison to years of experience with verbs as commands. Prior studies have shown only three dogs that consistently retrieved objects given a verbal referent (Kaminski et al., 2004; Pilley and Reid, 2011). Additionally, those dogs had been trained to retrieve from a young age (<11 months), and in most cases rarely attained 100 percent accuracy. Object retrieval training for the current experiment was modeled from these studies; however, because the dogs’ owners conducted training at home on a voluntary basis, training rigor could not be enforced.

Although humans readily generalize the meaning of words to a variety of contexts, this may not be the case for dogs. The environment in which the dogs learned the words was different than both the testing and scanning environments (Mills, 2015). In addition, although human fMRI language studies do not typically repeat the spoken word each trial, as is common in oddball paradigms, it was necessary for the dogs to make sure that they heard each word. Trials also did not include a condition in which a spoken pseudoword was followed by a trained object, or trials in which a trained object was mismatched to a trained word. These types of trials would have provided additional evidence for violation of expected semantic content; however, these types of trials have the potential to confuse the dogs and result in extinction of the words already learned. Lastly, dogs might have habituated to the continued presentation of trained words followed by trained objects, as opposed to the single trial presentations of pseudowords and the accompanying novel objects.

So what do words mean to dogs? Even though our findings suggest a prominent role for novelty in dogs’ processing of human words, this leaves the question of what the words represent. One possibility is that the words had no further representation than the relative hedonic value of the objects. While some dogs showed a behavioral preference for one object over the other, this preference was not reflected in whole brain analyses. Admittedly, the somewhat arbitrary designation of word1 / word2 and object1 / object2 could explain the nonsignificant results in the univariate analysis. Indeed, the MVPA of word1 vs. word2, which identified regions that classified the words above chance regardless of directionality, showed one cluster in the left caudate. However, the MVPA also identified clusters in the left TPJ, amygdala, and posterior thalamus. The TPJ was located just posterior to the region in the univariate analysis, which would take it out of the area of cortex associated with low-level acoustic processing. Its location appears similar to human angular gyrus – aka Wernicke’s area. If so, this could be a potential site for receptive word processing in dogs (e.g. the Dog Wernicke’s Area), but future work would need to verify this.

Evaluating classifier performance for MVPA remains a complex task. We used MVPA as an exploratory analysis to identify brain regions that potentially discriminate between trained words across dogs. But classification using the whole brain may result in a high classification accuracy that is not generalizable across subjects. Indeed, the regions identified using MVPA were of marginal statistical significance, especially given the small sample size. Further, it should be noted that only a subset of dogs contained informative voxels in the TPJ region. Although all dogs had informative voxels somewhere in the brain, only seven dogs had informative voxels in the TPJ area. Thus, even though all the dogs were cleared for scanning by reaching performance criterion, they may have used different mechanisms to process the words. Like our previous fMRI studies, heterogeneity seems to be the rule (Cook et al., 2016a; Cook et al., 2016b). Even so, the accuracy of the classifier was not correlated with a dog’s performance. This suggests that performance on such tasks may be influenced by factors other than word discrimination alone.

These results highlight potential mechanisms by which dogs process words. Word novelty appears to play an important role. The strong response of the parietotemporal region to pseudowords suggests that dogs have some basic ability to differentiate words with associations from those that do not. Future studies may reveal whether these representations remain in the auditory domain or whether such representations are invariant to modality.

## Acknowledgments

Thanks to Dr. Kate Revill for advice about language processing and generating pseudowords. Thank you to all of the owners who trained their dogs over the course of 6 months for this study: Lorrie Backer, Darlene Coyne, Vicki D’Amico, Diana Delatour, Marianne Ferraro, Patricia King, Cecilia Kurland, Claire Mancebo, Cathy Siler, Lisa Tallant, Nicole & Sairina Merino Tsui, and Nicole Zitron.

## Author Contributions

A.P., P.C., and G.B. designed and performed research; A.P, R.C., and G.B. analyzed data; A.P., P.C., G.B., & M. S. trained dogs; and A.P., G.B., P.C., R. C., and M.S. wrote the paper.

## References

Andics, A., Gabor, A., Gacsi, M., Farago, T., Szabo, D., and Miklosi, A. (2016). Neural mechanisms for lexical processing in dogs. Science 353, 1030–1032.

Andics, A., Gacsi, M., Farago, T., Kis, A., and Miklosi, A. (2014). Voice-sensitive regions in the dog and human brain are revealed by comparative fMRI. Curr Biol 24, 574–578.

Ann Young, C. (1991). Verbal commands as discriminative stimuli in domestic dogs (Canis familiaris). Applied Animal Behaviour Science 32, 75–89.

Avants, B.B., Tustison, N.J., Song, G., Cook, P.A., Klein, A., and Gee, J.C. (2011). A reproducible evaluation of ANTs similarity metric performance in brain image registration. Neuroimage 54, 2033–2044.

Bergelson, E., and Swingley, D. (2012). At 6-9 months, human infants know the meanings of many common nouns. Proc Natl Acad Sci U S A 109, 3253–3258.

Berns, G.S., Brooks, A., and Spivak, M. (2013). Replicability and heterogeneity of awake unrestrained canine FMRI responses. PLoS One 8, e81698.

Berns, G.S., Brooks, A.M., and Spivak, M. (2012). Functional MRI in awake unrestrained dogs. PLoS One 7, e38027.

Berns, G.S., Brooks, A.M., and Spivak, M. (2015). Scent of the familiar: An fMRI study of canine brain responses to familiar and unfamiliar human and dog odors. Behav Processes 110, 37–46.

Bloom, P. (2004). Behavior. Can a dog learn a word? Science 304, 1605–1606.

Brazdil, M., Dobsik, M., Mikl, M., Hlustik, P., Daniel, P., Pazourkova, M., Krupa, P., and Rektor, I. (2005). Combined event-related fMRI and intracerebral ERP study of an auditory oddball task. Neuroimage 26, 285–293.

Cacciaglia, R., Escera, C., Slabu, L., Grimm, S., Sanjuan, A., Ventura-Campos, N., and Avila, C. (2015). Involvement of the human midbrain and thalamus in auditory deviance detection. Neuropsychologia 68, 51–58.

Cook, P.F., Brooks, A., Spivak, M., and Berns, G.S. (2016a). Regional brain activations in awake unrestrained dogs. Journal of Veterinary Behavior-Clinical Applications and Research 16, 104–112.

Cook, P.F., Prichard, A., Spivak, M., and Berns, G.S. (2016b). Awake canine fMRI predicts dogs’ preference for praise vs food. Soc Cogn Affect Neurosci 11, 1853–1862.

Cook, P.F., Spivak, M., and Berns, G.S. (2014). One pair of hands is not like another: caudate BOLD response in dogs depends on signal source and canine temperament. PeerJ 2, e596.

Cox, R.W., Chen, G., Glen, D.R., Reynolds, R.C., and Taylor, P.A. (2017). FMRI Clustering in AFNI: False-Positive Rates Redux. Brain Connect 7, 152–171.

Cuaya, L.V., Hernandez-Perez, R., and Concha, L. (2016). Our Faces in the dog’s brain: Functional imaging reveals temporal cortex activation during perception of human faces. PLoS One 11, e0149431.

D’aniello, B., Scandurra, A., Alterisio, A., Valsecchi, P., and Prato-Previde, E. (2016). The importance of gestural communication: a study of human-dog communication using incongruent information. Anim Cogn 19, 1231–1235.

Datta, R., Lee, J., Duda, J., Avants, B.B., Vite, C.H., Tseng, B., Gee, J.C., Aguirre, G.D., and Aguirre, G.K. (2012). A digital atlas of the dog brain. PLoS One 7, e52140.

Dilks, D.D., Cook, P., Weiller, S.K., Berns, H.P., Spivak, M., and Berns, G.S. (2015). Awake fMRI reveals a specialized region in dog temporal cortex for face processing. PeerJ 3, e1115.

Eklund, A., Nichols, T.E., and Knutsson, H. (2016). Cluster failure: Why fMRI inferences for spatial extent have inflated false-positive rates. Proc Natl Acad Sci U S A 113, 7900– 7905.

Friederici, A.D., Meyer, M., and Von Cramon, D.Y. (2000). Auditory language comprehension: An event-related fMRI study on the processing of syntactic and lexical information. Brain Lang 74, 289–300.

Fukuzawa, M., Mills, D.S., and Cooper, J.J. (2005). The effect of human command phonetic characteristics on auditory cognition in dogs (Canis familiaris). J Comp Psychol 119, 117–120.

Goldman, R.I., Wei, C.Y., Philiastides, M.G., Gerson, A.D., Friedman, D., Brown, T.R., and Sajda, P. (2009). Single-trial discrimination for integrating simultaneous EEG and fMRI: identifying cortical areas contributing to trial-to-trial variability in the auditory oddball task. Neuroimage 47, 136–147.

Hanke, M., Halchenko, Y.O., Sederberg, P.B., Hanson, S.J., Haxby, J.V., and Pollmann, S. (2009). PyMVPA: A python toolbox for multivariate pattern analysis of fMRI data. Neuroinformatics 7, 37–53.

Howell, T.J., Conduit, R., Toukhsati, S., and Bennett, P. (2012). Auditory stimulus discrimination recorded in dogs, as indicated by mismatch negativity (MMN). Behav Processes 89, 8–13.

Humphries, C., Binder, J.R., Medler, D.A., and Liebenthal, E. (2006). Syntactic and semantic modulation of neural activity during auditory sentence comprehension. J Cogn Neurosci 18, 665–679.

Kaminski, J., Call, J., and Fischer, J. (2004). Word learning in a domestic dog: evidence for “fast mapping”. Science 304, 1682–1683.

Keuleers, E., and Brysbaert, M. (2010). Wuggy: A multilingual pseudoword generator. Behav Res Methods 42, 627–633.

Kiehl, K.A., Laurens, K.R., Duty, T.L., Forster, B.B., and Liddle, P.F. (2001). An event-related fMRI study of visual and auditory oddball tasks. Journal of Psychophysiology 15, 221–240.

Kotz, S. (2002). Modulation of the lexical–semantic network by auditory semantic priming: An event-related functional MRI study. NeuroImage 17, 1761–1772.

Linden, D.E., Prvulovic, D., Formisano, E., Vollinger, M., Zanella, F.E., Goebel, R., and Dierks, T. (1999). The functional neuroanatomy of target detection: an fMRI study of visual and auditory oddball tasks. Cereb Cortex 9, 815–823.

Mahmoudi, A., Takerkart, S., Regragui, F., Boussaoud, D., and Brovelli, A. (2012). Multivoxel pattern analysis for FMRI data: A review. Comput Math Methods Med 2012, 961257.

Markman, E.M., and Abelev, M. (2004). Word learning in dogs? Trends Cogn Sci 8, 479–481; discussion 481.

Mills, D.S. (2015). What’s in a word? A review of the attributes of a command affecting the performance of pet dogs. Anthrozoös 18, 208–221.

Misaki, M., Kim, Y., Bandettini, P.A., and Kriegeskorte, N. (2010). Comparison of multivariate classifiers and response normalizations for pattern-information fMRI. Neuroimage 53, 103–118.

Müller, C.A., Schmitt, K., Barber, A.L.A., and Huber, L. (2015). Dogs can discriminate emotional expressions of human faces. Current Biology 25, 601–605.

Persson, M.E., Roth, L.S., Johnsson, M., Wright, D., and Jensen, P. (2015). Human-directed social behaviour in dogs shows significant heritability. Genes Brain Behav 14, 337–344.

Pilley, J.W., and Reid, A.K. (2011). Border collie comprehends object names as verbal referents. Behav Processes 86, 184–195.

Raettig, T., and Kotz, S.A. (2008). Auditory processing of different types of pseudo-words: an event-related fMRI study. Neuroimage 39, 1420–1428.

Slotnick, S.D. (2017). Cluster success: fMRI inferences for spatial extent have acceptable false-positive rates. Cogn Neurosci 8, 150–155.

Stelzer, J., Chen, Y., and Turner, R. (2013). Statistical inference and multiple testing correction in classification-based multi-voxel pattern analysis (MVPA): random permutations and cluster size control. Neuroimage 65, 69–82.

Varoquaux, G. (2017). Cross-validation failure: Small sample sizes lead to large error bars. Neuroimage.

Waxman, S., Fu, X., Arunachalam, S., Leddon, E., Geraghty, K., and Song, H.J. (2013). Are nouns learned before verbs? Infants provide insight into a longstanding debate. Child Dev Perspect 7, 155–159.

Zaine, I., Domeniconi, C., and Costa, A.R.A. (2014). Exclusion performance in visual simple discrimination in dogs (Canis familiaris). Psychology & Neuroscience 7, 199–206.

